# Polarized neutrons for the study of individual and collective fast dynamics in proteins

**DOI:** 10.64898/2026.08.30.748099

**Authors:** Agathe Nidriche, Jacques Ollivier, J Ross Stewart, Judith Peters

## Abstract

Neutron scattering is a powerful technique to investigate atomic structures and molecular dynamics of proteins at the nano-scale. When it comes to dynamics, incoherent and coherent scattering respectively provide information on the single and collective dynamics of nuclei. In proteins, hydrogen has the highest incoherent cross-section, and it is common practice to overlook the contribution of coherent terms stemming from all nuclei. However, the fast collective dynamics of heavier nuclei could also be studied if coherent scattering and incoherent scattering were experimentally separated. The recent advent of polarized neutron spectroscopy with sufficient flux and energy resolution has made it possible, and opens new perspectives to investigate the relative importance of coherent scattering and the information it provides on biological samples. The present study reports on the use of polarized quasi-elastic neutron scattering (QENS) and the application of a minimalistic model adapted to both individual and collective dynamics. Using a perdeuterated green fluorescent protein as a model globular protein, the study provides an interpretation of the dynamical parameters obtained with QENS, and a comparative study of the Elastic Coherent and Incoherent Scattering Factor. Based on both experiments and calculations, we discuss the relative importance of distinct and self components of coherent scattering, which is often wrongly assumed to be representative of collective dynamics only. The results highlight the current impediments rendering complicated a straightforward analysis of fast collective dynamics in hydrated protein samples.

## I. INTRODUCTION

The dynamics of proteins is characterized by a vast spectrum of timescales, ranging from femtoseconds to hours, where fast internal diffusive motions in the picosecond regime are often fundamental to initiate slower functional dynamics. [1] Thermal neutron scattering is an ideal technique for studying the structural dynamics of condensed matter at the atomic scale, since thermal neutrons have the unique property that their wavelengths are comparable with typical interatomic distances, while their energies are comparable to the energies of the nuclei in the scattering system. The measured scattering intensities give information about correlated atomic motions in space and time, which are contained in the dynamic structure factor, *S*(**q**, *ω*). Its arguments, **q** and *ω*, are, respectively, the momentum and energy transfer from the neutron to the sample (in units of *ħ*) and inversely related to the space- and time-scale. The dynamic structure factor can be split into a coherent part, reflecting collective motions in the scattering system, and an incoherent part depicting the system-averaged single atom dynamics. Each nucleus has a coherent and an incoherent cross section for neutron scattering, which are isotope-dependent [2]. At fixed momentum transfer a neutron scattering spectrum can be schematically decomposed into an elastic part, corresponding to immobile scatterers and motional amplitudes reflecting the geometry of relaxations, a quasi-elastic part, representing diffusive motions and relaxation processes, and an inelastic part, reflecting vibrational motions with well-defined frequencies. Noting that incoherent scattering from hydrogen is largely dominant and that biomolecular systems contain a large number of hydrogen atoms (≈ 50 %), neutron scattering studies of such systems are traditionally performed on samples in which the protonated biomolecules are immersed in a deuterated solvent, considering either hydrated powders or aqueous solutions. The assumption is here that the contribution from the solvent can be neglected due to the lower incoherent scattering cross section of deuterium, and only incoherent scattering from the hydrogen atoms in the solute molecule is seen [2, 3]. Coherent contributions become important when a protonated sample is dissolved in a deuterated buffer or when both the sample and its environment are perdeuterated. In a perdeuterated hydrated protein, all nuclei participate with similar magnitudes to the coherent scattering intensity, in contrast to the incoherent intensity which is dominated by hydrogen isotopes. In addition, the contributions of the coherent and incoherent parts depend on the momentum transfer **q** between the neutron and the scattering atom, thus their relative parts can vary [4]. Recently, there have been extensive discussions in the community aimed at understanding whether the addition of polarization analysis to new, or existing instruments would allow access to more interesting results including individual and fast collective dynamics measured on the same sample at the cost of beam intensity. As only few studies were published on concrete examples, we wanted to contribute with a systematic investigation and show what is feasible and what kind of information can be obtained. It is up to the interested researcher to judge what is useful, but this is only possible when knowing the potential implications of using polarization analysis.

In the present study, we investigate quasi-elastic neutron scattering (QENS) from a model protein containing only a very small amount of hydrogen atoms: it is a perdeuterated sample of green fluorescent protein (GFP) in powder form and hydrated at a level of 0.4 g D_2_O/g sample. GFP is indeed one of the few proteins which can be produced in both perdeuterated and protonated forms. The choice of a hydrated powder form was motivated by the fact that powders provide a system widely adopted in investigations at temperatures below the water freezing point that permits the observation of several interesting phenomena: rehydrated powders can retain the folded structure of proteins and mimic their native confined environment. Global diffusion of the whole protein is suppressed in powder forms, but here we are studying distinct coherent contributions as collective motions.

We measured the same sample on two time-of-flight (TOF) spectrometers: IN5 [5] at the Institut Laue Langevin (ILL) in Grenoble, France, and on LET [6, 7] (ISIS Neutron and Muon Source, UK) using uniaxial polarization analysis. This is so far one of the few instruments that permits the separation of the coherent and incoherent parts of *S(***q**, *ω)*. Otherwise the two instruments can be run with very similar characteristics (in terms of instrumental resolution and *q*-range) to justify the direct comparison of the outputs with the aim to identify the impact of the coherent part on the data.

Traditionally, polarization analysis was used for investigations of magnetic materials [8], in order to decipher complex magnetically ordered structures, or to separate magnetic scattering from nuclear scattering. Polarization analysis provides four different projections of the dynamic structure factor, which are averaged in the case of unpolarized neutrons. Nuclei in biomaterials do not present spin ordering, since they are supposed to be randomly oriented. Therefore, only two projections of *S(***q**, *ω)* are accessible, which are linearly combined to obtain the incoherent and coherent parts of scattering. This presents a powerful tool to isolate self and collective dynamics in biological samples. Quite early, soft matter was investigated with polarized neutrons by Gabrys, Zajac, Schärpf et al. [9–12]. They pointed out the relevance of the technique and already started to tackle coherent contamination in elastic and quasi-elastic neutron scattering. Later, Gaspar et al. [13] measured polarized diffraction spectra for different types of protein powders to get access to a broad *q*-range. However, their measurements were limited to static structure factors *S(***q)**. The results showed the expected differences between the coherent and incoherent parts, but also a strong dependence on the sample hydration level. Here we expand polarization analysis studies to dynamical investigations on proteins and compare the results directly to the outcome from IN5, which gives access to the total signal, only.

In the present work, we further analyse the data from both instruments with the same minimalist approach developed by G. Kneller [14] and show that it is suitable for the description of coherent and incoherent parts of neutron scattering, and for comparison between similar instruments. We also investigate the impact of coherent dynamics on elastic scattering temperature scans, which are often performed to provide information on average hydrogen mean-squared displacements. Finally, we discuss the question of whether or not it is necessary to perform polarization analysis for all standard studies of hydrogen-rich biological samples contrasted by D_2_O, as well as which topics in protein dynamics would grandly benefit from using this technique for QENS studies.

## II. NEUTRON SCATTERING

### A. Scattering functions

The probed quantity in a scattering experiment with thermal neutrons is the differential scattering cross section,

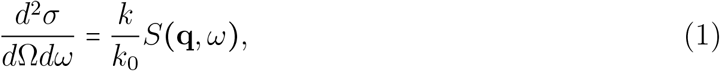

where *k* et *k*_0_ are the momenta of the scattered and incoming neutrons, respectively, in units of *ħ* and the dynamic structure factor, *S*(**q**, *ω*), is the quantity of interest which carries the information about the structural dynamics in the scattering system. As any spectroscopic technique, neutron scattering probes the Fourier transform of specific time correlation functions. Considering a scattering experiment with unpolarized neutrons, the dynamic structure factor has the form

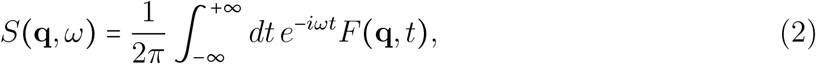

where the intermediate scattering function, F(**q**, *t*), is the sum of a coherent component, containing information about the structural dynamics of the atoms in the system and from cross-terms between the different atoms, and an incoherent component, informing about the system-averaged dynamics of single atoms only,

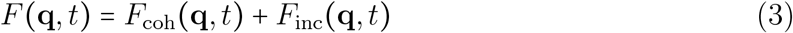

with

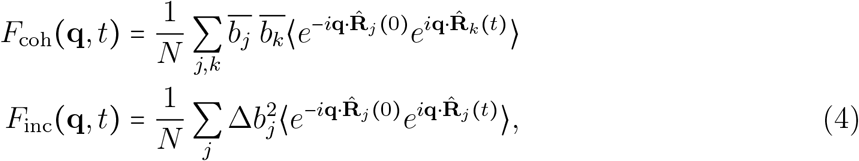

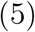

where 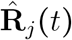 is the position operator of atom *j* (precisely of its nucleus) in the Heisenberg picture. The symbol 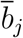 stands for the mean scattering length [2, 15] of atom *j* and 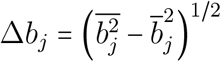 is the corresponding standard deviation. The mean is here taken over all isotopes and over the relative orientations of nuclear and neutron spin at the moment of interaction.

### B. Polarization analysis

Neutrons have a spin of 1/2 and therefore can align either parallel or antiparallel to an applied magnetic field with angular momenta *s*_*z*_ = ± *ħ/*2, giving spin-up (∣↑⟩) and spin-down (∣↓⟩) eigenstates. The neutron beam polarization is given by the normalised difference between the numbers of neutrons in each of these states, 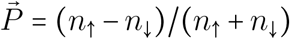. In scattering experiments with neutron polarization analysis, the polarization of the incoming neutrons is prepared in one of the possible states and the scattering intensities are recorded separately for neutrons without spin flip (⇈) and with spin flip (↑↓). To account for this additional information, the calculation of the differential scattering cross section must be changed correspondingly by fixing the initial and final polarization of the neutrons, instead of summing over them. The corresponding dynamic structure factors can be expressed by the coherent and incoherent dynamic structure factors obtained from scattering with unpolarized neutrons [16],

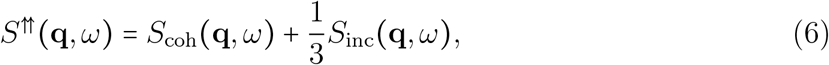

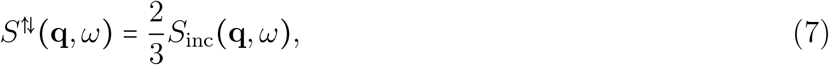

and these relations may then be inverted to yield explicit expressions for the coherent and incoherent components of the dynamic structure factors,

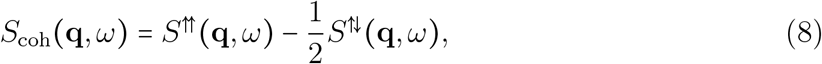

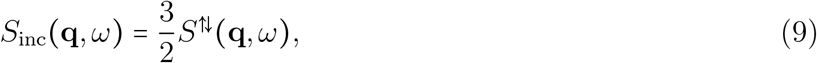

which are superposed in neutron scattering spectra obtained without polarization analysis.

### C. Modelling of *F*(*q, t*)

In order to analyze the QENS data, a modified version of Schofield’s semiclassical approximation is applied to *F*(**q**, *t*) [17] which preserves the correct normalization of the scattering functions when working with classical mechanics and which has been successfully used in earlier work [18–21]. Scattering of protein powders is isotropic, therefore the scattering functions depend on the modulus *q* ≡ |**q**| of the momentum transfer vector, only.

Subsequent steps consist of defining which variables are implied in the modelling of incoherent and coherent scattering, and introducing a model for the relaxation of those variables in the case study of complex amorphous materials.

#### 1. Generic form for the intermediate scattering function

Following ref. [20], the intermediate function *F*(*q, t*) defined in equation 4 is expressed as the autocorrelation of a time-dependent dynamical variable *ρ*_*q*_ (*t*)

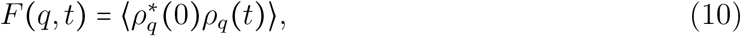

or alternatively as the relaxation of a function *ϕ*_*q*_ (*t*) depending on *ρ*_*q*_ (*t*), split into static and dynamic components corresponding respectively as elastic and inelastic scattering:

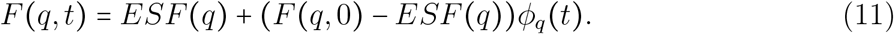

The relaxing variable is *δρ*_*q*_ (*t*), the deviation of *ρ*_*q*_ (*t*) from equilibrium

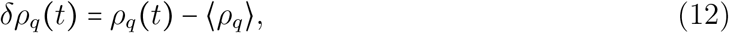

such that *ϕ (t)* is the relaxation function driving the evolution of the dynamical variable

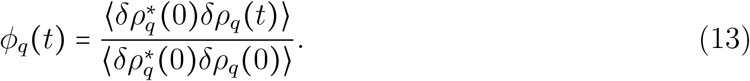

This implies that the intermediate scattering function *F*(*q, t*) relaxes from an initial state, the static structure factor *S*(*q*)

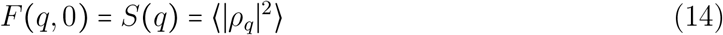

which equals 1 in the case of incoherent scattering, to its asymptotic value in time, the elastic scattering factor *ESF* (*q*),

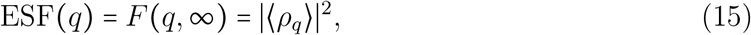

which is the weighted sum of the elastic coherent scattering factor ECSF (*q)* and the elastic incoherent scattering factor EISF(*q*).

Now, separating the intermediate scattering function into its incoherent and coherent components, one finds that the evolution of the system is driven by the sum of the relaxation of two variables, *δρ*_q,inc_ (*t)* and *δρ*_q,coh_ (*t)* .

The incoherent dynamical variable *δρ*_q,inc_ (*t)* involves the position *x* of a single atom which is representative of the average nucleus in the sample. *x* is here the projection of the position operator 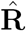. In proteins, the abundance of hydrogen H and its important relative cross-section makes it the most representative nucleus in incoherent scattering:

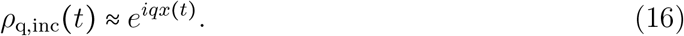

The coherent dynamics variable *δρ*_q,coh_ (*t)* involves the weighted sum of all different nuclei *k*, providing structural information:

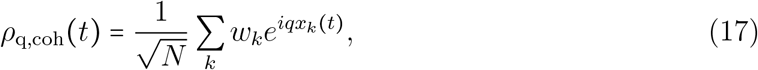

where *w*_*k*_ are weights that are normalized in order to recover *ϕ*_*q*_(0) = 1.

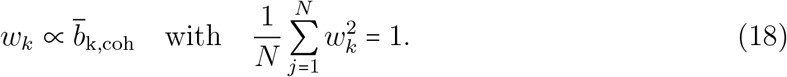

Therefore, we arrive at a general expression for the dynamical variable *δρ*_*q*_:

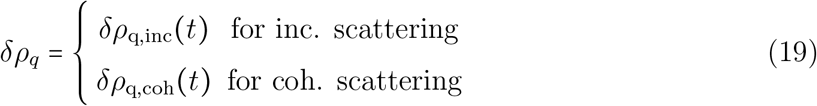

#### 2. A self similar model accounting for both coherent and incoherent scattering

The choice of *ϕ (t)* guides the physical interpretation. To describe the internal relaxational dynamics of proteins, the approach from ref [20] introducing a fractional Ornstein Uhlenbeck (fOU) model [22, 23] depicting non-equilibrium processes in a rugged harmonic potential is used [18, 24, 25]. In this frame, the relaxation function is the following

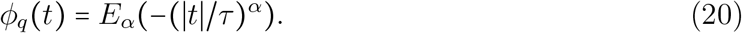

Here 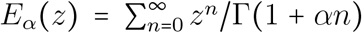 denotes the Mittag-Leffler (ML) function [26, 27], a generalised exponential function where *τ* ≡ *τ* (q) sets a time scale and *α* ≡ *α* (q) is a form parameter.

In common with other biological entities, proteins are reported to follow self-similar dynamics over large ranges of timescales, from the very fast localised diffusive motions at the ps-scale to the longer timescales of biological functions (> *s*) [28, 29]. The Mittag-Leffler function bears significant mathematical properties describing well the relaxation properties required here :

- The Mittag-Leffler function is the solution of the fractional stochastic diffusion in a harmonic potential [25], which describes local small amplitude motions in a rugged potential analogous to the energy landscape of a protein [30].
- It tends to a power law at long times: 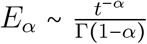. Therefore, it is asymptotically a self-similar function and is form-invariant under tranformation *t* → *µ t* for any *µ >* 0. Furthermore, any relaxation that follows asymptotically a power law decay and for which the memory kernel reaches its asymptotic form almost instantaneously follows a Mittag-Leffler function [31].
- The Mittag Leffler function interpolates between a stretched exponential for time *t* → 0^+^ (an exponential function is retrieved for *α* → 1) and a power law function for time *t* → ∞ [32], therefore embeds all the models commonly used in QENS. This choice of model is adapted to the study of proteins which undergo simultaneously many relaxation processes that cannot be discriminated using simple exponential decays [33].

#### 3. Interpretation of parameters

In such a way, *F*(*q, t*) is parametrized with 3 *q*-dependent parameters:

*α*, **the form parameter**. *α* ∈ [0, 1] yields a mono-exponential *ϕ (t)* function for *α =* 1 and illustrates the heterogeneity of internal processes in the protein. It can be understood either as a distribution of exponential decays with a broad range of relaxation times or as the non-exponential decay of individual nuclei.

In the case of multi-exponential decay [25] it implies that

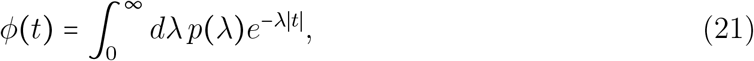

where *p*(*λ*) is a positive normalized function of the form

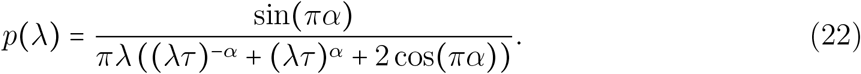

Using the result of a seminal paper by Zwanzig on the calculation of effective short time diffusion constants for diffusion in a rough potential [34] and a harmonic form of the envelope potential, the distribution of relaxation rates can be converted into a distribution of energy barriers [20, 35],

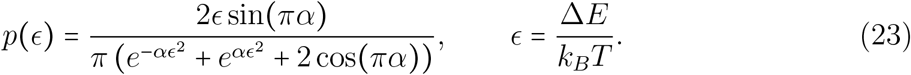

Here Δ*E* is the energy difference between the rough potential and the smooth harmonic envelope potential. In that case, the rugged potential in which the dynamical variable evolves follows

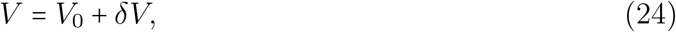

where

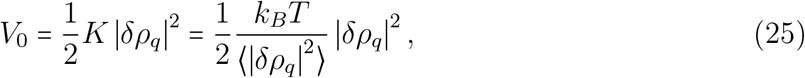

is parametrized by ⟨∣*δρ*_*q*_∣^2^⟩, the Mean-Square atomic Position Fluctuations (MSPFs) of the dynamical variable. *δV* is distributed following a Gaussian distribution of standard deviation *ϵ*, and is therefore also dependent on *δρ*_*q*_.

*τ*, **the scale factor**, sets the time scale at which dynamics occur, which is highly dependent on the instrumental resolution. More precisely, it parametrizes *p(λ)* (equation 22) which is the rates distribution. *P(λ)* already broadens strongly around 1/*τ* for *α =* 0.99. With decreasing *α*, its peak value shifts down and diverges at *λ* = 0 when *α* → 0, implying the prevalence of extremely long time scales.

**ESF (***q)*, **the asymptotic value of the function** *F(q, t)* **for** *t* → ∞. ECSF(*q)* is scarcely mentioned in literature for the study of proteins [11, 36], either due to the assumption that it is negligible or to the requirement for deuterated compounds to extract the quantity without the use of polarized neutrons.

Conversely, EISF(*q)* is a quantity which is central to study the geometry of diffusion in the frame of standard models [37]. It is fitted with the diffusion model and takes into account the dynamics of the system in the range of study. The EISF is also ubiquitously measured as a function of temperature for proteins in powder form since it provides a fast and simple understanding of the flexibility of proteins through the average mean square displacements of H atoms (MSDs) [38]. However, those standard measurements of temperature-dependent MSDs are based on short time-window measurements in the hypothesis of a dominant elastic peak, which is a model free approach. Therefore, in this case the diffusion contained in the resolution function of the instrument is treated as elastic, as detailed by Kneller *et al*. [39]. The notation further used for this quantity obtained using the elastic fixed window method is denoted EISF_*m*_*(q)* .

The elastic fixed window method places itself in the assumption of small *q*-values and isotropic scattering. We introduce Δ*x*, the deviations of the position of the representative nucleus from its mean value, corresponding to Mean-Square Position Fluctuations (MSPF)

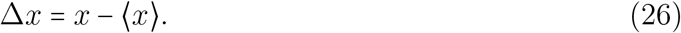

The cumulant expansion of intermediate scattering function [40] yields a Gaussian form for the EISF_*m*_, which is the quantity measured in elastic fixed window scans

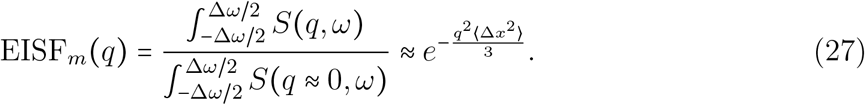

In the limiting case of homogeneity in the atomic motions, the MSPFs converge towards the MSDs [41], which are obtained from the Gaussian approximation [42].

Since for small *q*-values the static quantities ⟨∣*ρ*_*q*_∣^2^⟩, and ⟨∣*ρ*_*q*_∣^2^⟩ can be related to the position fluctuations of the atoms in the scattering system, we use the MSPFs measured with the EISF as a proxy for the quantity ⟨∣*δρ*_*q*_∣^2^⟩ for the sketch of the rugged potential, equation 24.

## III. MATERIALS AND METHODS

### A. Sample preparation

A perdeuterated sample of green fluorescent protein (GFP) of atomic formula H_1505_D_414_C_1261_N_342_O_379_S_8_ of about 100 mg dry mass was prepared by the Deuteration Lab, at Institut Laue Langevin (ILL), France. We refer to this deuterated protein as “dGFP”. The preparation is detailed in Ref. [21] (Supplementals). It was first lyophilised and then re-hydrated to a level of *h* = 0.4 with *h* = g protein/ g D_2_O in a desiccator with D_2_O filled atmosphere. Mass spectrometry revealed over 99% of deuteration of non-labile atoms.

### B. Experiments

#### 1. Quasi-elastic neutron scattering

Data analysis is performed following the procedure described in Refs [18–21]. QENS data were acquired with the two Time-of-Flight (TOF) spectrometers IN5 (ILL, DOI Ref. [43]) [5] and LET (ISIS Neutron and Muon Source, UK, DOI Ref. [44]) [45, 46] with similar characteristics, summarized in Table I. While LET is equipped with polarization analysis, IN5 uses an unpolarized beam. Uniaxial polarization analysis on LET consists of polarizing the neutron beam by means of a supermirror polarizer, and modifying the projection of the spin with a precession coil flipper and a field ramp synchronised to the ISIS pulse [47]. The scattered neutron polarization is analyzed with a hyperpolarized ^3^He neutron spin filter. The neutrons are then detected by an array of position sensitive high-pressure ^3^He counters.

**TABLE I.** Measurement setups on IN5 TOF and LET TOF.

|  | IN5 | LET |
| --- | --- | --- |
| Polarization analysis setup | Not equipped | Equipped |
| Neutron source | Continuous | Pulsed |
| Incident neutron energy [meV] | 3.27 | 3.27 |
| Energy resolution $\delta E$ [$\mu\text{eV}$] | 77 | 95 |
| $t_{\text{max}}$ [ps] | 16 | 28.5 |
| $\Delta t$ ($\equiv t_{\text{min}}$) [ps] | 1.2 | 1.2 |
| $q$ [$\text{\AA}^{-1}$] | [0.8, 2] | [0.6, 2.2] |
| $\Delta q$ [$\text{\AA}^{-1}$] | 0.1 | 0.2 |

In order to compare the dynamical structure factors obtained with IN5 and LET, we sum incoherent and coherent contributions on LET to yield the total scattering dynamic structure factor,

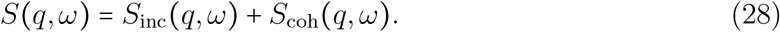

The QENS spectra from the powder samples were taken at the physiological temperature *T =* 310 K, to provide the measured dynamical structure factor *S(q, ω)*, and the data from the sample at *T =* 2 K were used to define the resolution function of the instrument. Measurements with a vanadium sample were performed to correct for differences in detector efficiency, and with an empty aluminium cell to subtract the scattering from the sample holder, which was flat in case of IN5 (thickness *t =* 0.6mm) and annular cylinder-shaped in case of LET, *t =* 1mm, diameter *d =* 16mm, height *h =* 40mm). The background signal on LET was subtracted using a run with cadmium [48].

We measured multiple incident energies on LET, using repetition-rate-multiplication [45]. We optimized the chopper frequencies for *E*_*i*_ *=* 3.27 meV to match IN5 characteristics. This resulted in low neutron flux at the higher resolution *E*_*i*_ *=* 1.83 meV repetition and therefore data at this *E*_*i*_ could not be analyzed. Data analysis has been performed for the intermediate scattering functions as described in previous work [18–21].

The sample’s transmission *T* was estimated by calculations (*T*_calc_ ≈ 0.95), and further confirmed by measurements on D7 (ILL) using a flat sample holder, in normal incidence to the beam, to be equal to *T =* 0.974. Therefore, geometry-dependent absorption is supposed to be negligible. The data was corrected for absorption from flat sample on IN5 using well-known analytic estimations (LAMP software [49]) assuming an incidence angle of 135^°^ and a thickness of 0.6 mm. The thin sample also results in a negligible amount of multiple-scattering, which otherwise would swap spin-flip and non-spin flip scattering events in case of polarised neutrons, resulting in an increase of apparent coherent scattering [50].

#### 2. Elastic scattering

On IN5, elastic scattering data were obtained in the form of 5 minute scans measured for *N* = 234 temperature points in the range *T* ∈ [14, 310] K, and integrated over the fixed elastic window which corresponds to the FWHM of the resolution function *S*_*R*_*(q, ω)*.

On LET, elastic scattering data were acquired during the temperature ramp of the cryostat from 2K to 310K with alternating spin-flip (sf) and non spin-flip (nsf) conditions, then integrated over the fixed elastic window and summed over approximately 12 minutes for each of the 9 data points. Incoherent scattering was obtained from spin-flip scattering only, and coherent scattering from the difference of intensities corresponding, respectively, to neutrons with and without spin-flip (equations 9 and 8). The average temperature over the elastic window scan is derived accordingly for incoherent and coherent scattering.

## IV. RESULTS

### A. A minimalistic model to extract individual and collective dynamic in proteins

The minimalistic model is robust, as shown by its ablity to capture the same dynamics on two similar instruments (see Table I), despite the low signal-to-noise ratio of the polarized setup. This is shown by comparing the “total” scattering data collected on the two instruments, which is the sum of incoherent and coherent dynamic structure factors in the case of LET (see equation 28) and the standard dynamic structure factor in the case of IN5. Figures 1 a) and 1 b) show respectively the intermediate scattering function *F(q, t)* and the dynamic structure factor *S(q, ω)*. Due to slightly differing resolutions and different sample holder geometries, *F(q, t)* and *S(q, ω)* differ more and more for increasing *q* on both instruments, especially at short times (high *ω* values). However, plotting the corresponding fit parameters *τ, α* and ESF on Figure 2 clarifies that dynamical parameters *τ*_tot_ and *α*_tot_ are highly comparable, either obtained on IN5 (black markers) or LET (gray markers). Conversely, the ESF differs for the lowest and highest *q*-values, most probably due to differences in instrumental resolutions. Therefore, this model is sufficient to raise comparable analysis whether data is acquired under high neutron flux conditions (IN5) or using polarized neutrons setups for which the flux is reduced up to 10 times (LET).

**FIG. 1.**
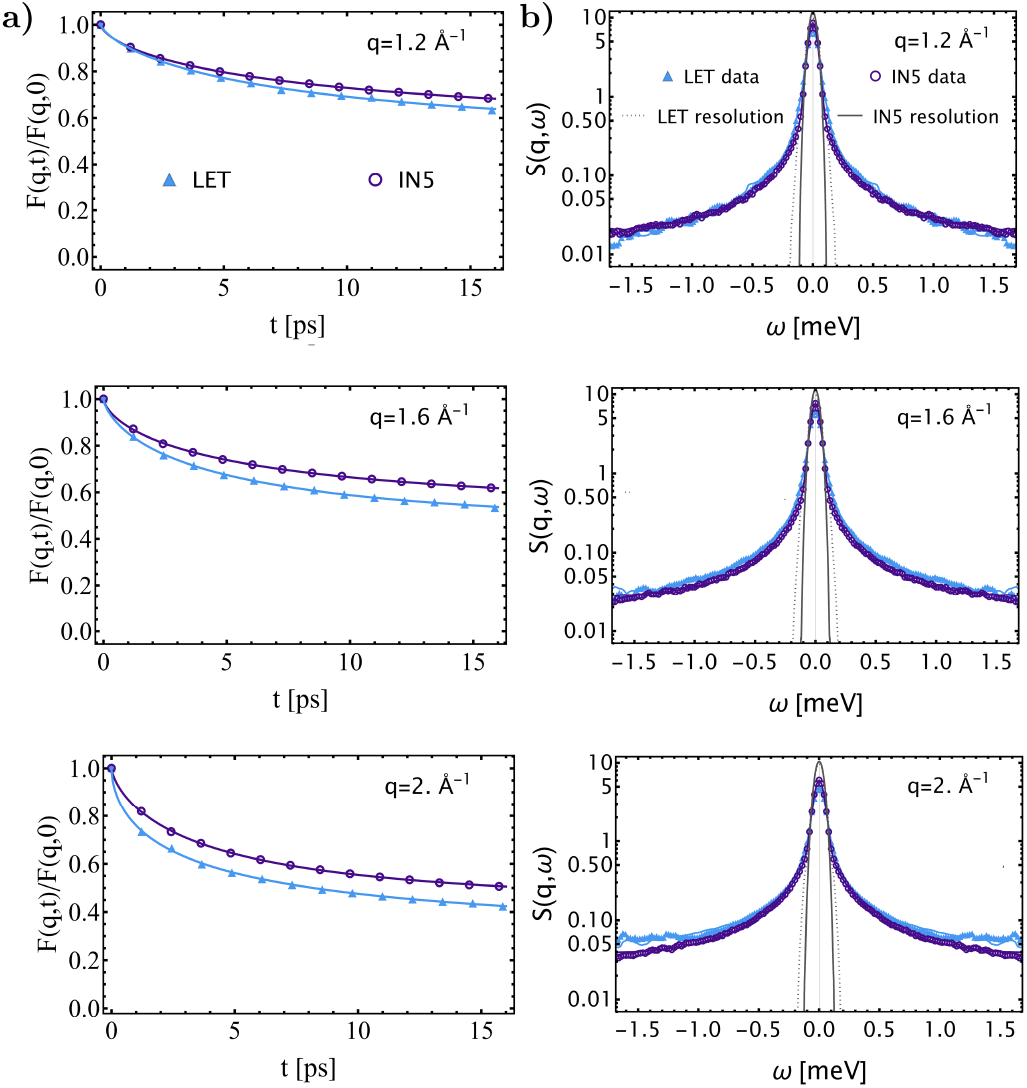
Scattering functions corresponding to total scattering measured at *T =* 310 K are displayed in light blue filled Δ markers for LET, equipped with polarization analysis, and in purple open ° marker for IN5, not equipped with polarization. Three different figures diplay increasing momentum transfers *q* ∈ *{*1.2, 1.6, 2}.Å^−1^. a) *F(q, t)*, intermediate function of dGFP hydrated powder. Full lines correspond to the model, equation 11. b) *S(q, ω)*, dynamic structure factor. Full lines correspond to the discrete Fourier Transform of the model function. The resolution of the instrument, obtained using the protein sample at *T =* 2K, is represented in black dotted lines for LET and in black full lines for IN5.

**FIG. 2.**
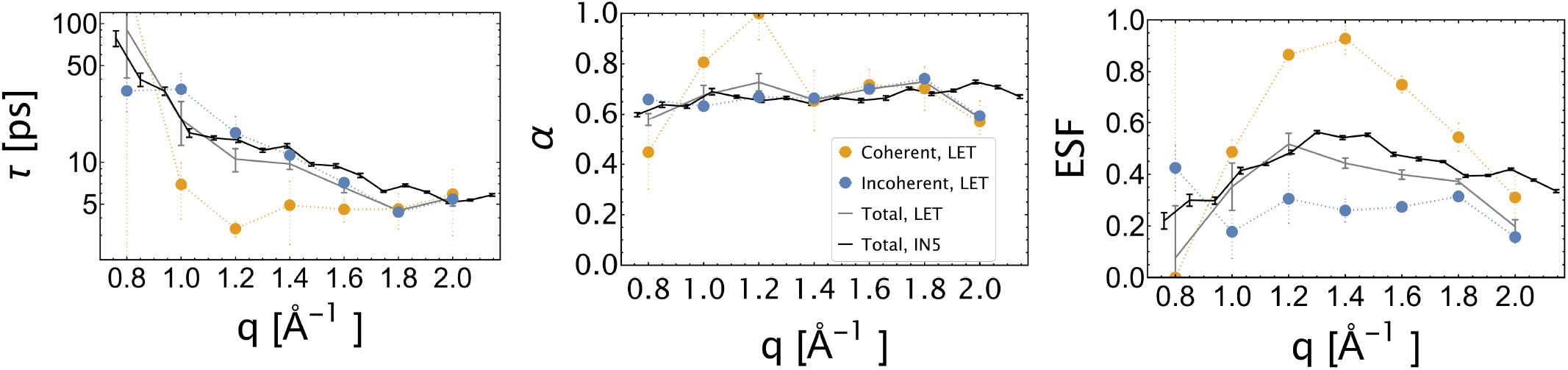
*q*-dependent total scattering dynamical parameters, *τ* on a semi-log scale, *α* and ESF, are compared for LET (gray markers), and IN5 (black markers), where joined lines are a guide for the eyes. Incoherent and coherent scattering parameters, obtained with LET, are displayed with filled blue and yellow markers, respectively.

Furthermore, this model is adapted to the study of both incoherent and coherent scattering: however, the diffusing dynamical variable is different in both cases, and captures respectively the average dynamics of a single nucleus or of an assembly of nuclei. Figure 2 compares the fit parameters obtained from total scattering, to the coherent and incoherent contributions that are extracted from LET data (filled yellow and blue markers, respectively). They show clear differences, proving the potential of the theoretical model to treat both polarized and non-polarized data. Local collective dynamics (*τ*_coh_) are decaying faster with respect to self-dynamics (*τ*_inc_) with differences above two sigma in the interval between 1 and 1.4 Å^−1^. The form parameter *α* is similar for incoherent and coherent scattering, and is of the order of magnitude of exponents found for complex amorphous materials. However, this parameter usually observes very slight changes even under modifications of external conditions, as for instance in the case of high hydrostatic pressure [51]. The motional amplitudes of the collective variable are higher than for the single nucleus variable, as observed with the comparison of the ECSF to the EISF. The *q*-dependence of collective motions, as indicated by *τ*_coh_ and ECSF, reflects the local structure of the protein and its hydration water, and show fast local collective motions from deuterated hydrogens comprised in the hydration layer. Conversely, *τ*_inc_ indicates the slow diffusive-like behaviour of the protein’s hydrogens. For a thorough discussion of the physical interpretation in terms of hydration water and internal protein motions, see Ref. [21].

Let us notice that, as discussed further in section V, the parameters extracted from total and incoherent scattering remain highly similar, despite the faster dynamics contained in coherent scattering. It can be explained by the relative weights of coherent and incoherent scattering functions (see section IV D), due to the large incoherent scattering length of hydrogen which dominates the other contributions.

Thus, despite seemingly different data upon visual inspection, the model enables high correspondence between instruments of similar characteristics: it makes it possible to compare the physical properties of a sample measured with and without polarization analysis setups. Furthermore, the same set of parameters can be used to describe coherent and incoherent scattering.

### B. Energy landscape interpretation in the frame of the minimalistic model

In a second step, the model provides an interpretation of data based on the energy landscape of proteins, and describes the diffusion of the dynamical variable in a rough harmonic potential. When a system diffuses in a harmonic potential, 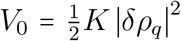, it undergoes a mono-exponential relaxation until equilibrium is reached. The non-equilibrium dynamics of folded proteins can, on the contrary, be considered as a fictitious particle evolving in a rugged energy potential with numerous free energy wells of height *δV* [35] in an harmonic envelope. The rough energy potential *V*_0_ + *δV* in which the variable diffuses is shown in Figure 3 a).

**FIG. 3.**
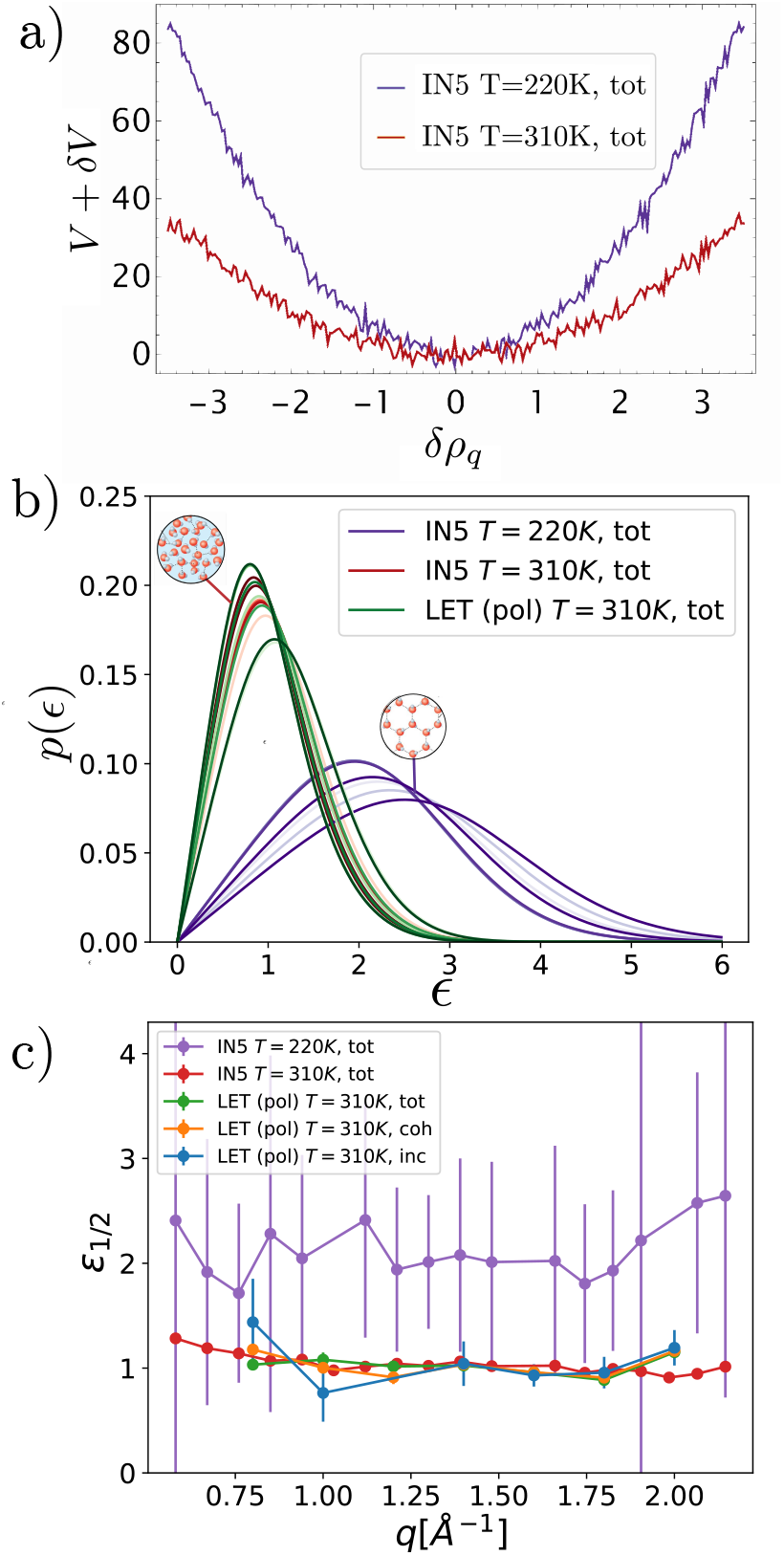
Study of energy barriers performed with IN5 and LET. Top panel: a) Sketch of the dimensionless potential *V*_0_ + *δV* as a function of *δρ*_*q*_ calculated for IN5 for *q* = 1.0 Å^−1^ . (IN5, *T =* 220 K: purple; IN5, *T =* 310 K: red). b) Unpolarized scattering distribution of dimensionless energy barriers *p(ϵ)* (equation 23) as a function of 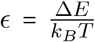, compared for the two instruments. The gradient of colours (IN5, *T* = 220 K: purple; IN5, *T* = 310 K: red; LET, *T* = 310 K: green) corresponds to a increase from *q* = 0.8 Å^−1^ to *q* = 2 Å^−1^ by steps of 0.2 Å^−1^. Sketches for Low Density Liquid (LDL) and High Density Liquid (HDL) phases of water are adapted from Ref [52]. (c) The median value of the energy barrier density, *ϵ*_1 /2_, is represented as a function of *q. ϵ*_1 /2_ as obtained for incoherent and coherent scattering (LET for *T* 310 K) are pictured in blue and orange, respectively.

To begin with, unpolarized data obtained on IN5 at *T =* 220 *K* and *T =* 310 *K*, corresponding respectively to the dynamical transition [53, 54] and the physiological temperatures, is used to illustrate the information provided by parameter *α* and its interpretation in terms of potential roughness. On the sketch provided in Figure 3 a), the harmonic envelope of the potential was obtained according to reference [35], where MSDs are used to estimate the force constant *K*. The potential gets softer when increasing temperature from *T =* 220K (in purple) to *T* = 310 K (in red). The fluctuations *δV* are given by a distribution of the energy barriers 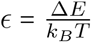 determined by equation 22.

Figure 3 b) shows a large shift of the density of energy barriers towards lower values from *T =* 220K to *T =* 310 K, which is better evaluated as a function of *q* introducing the median of the distribution, *ϵ*_1_/_2_, defined by 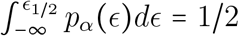, which is shown in Figure 3 c). *ϵ*_1_/_2_ expresses the capacity of the system to overcome transient energy barriers at given temperature conditions. It indicates a radical change of the local energy landscape experienced with temperature. LET total scattering data are in complete agreement for *T =* 310 *K*. Since the dynamics of the sample are dominated by hydration water, more information in Ref. [21], we interpret this in the frame of the liquid-liquid phase transition in supercooled hydration water around *T =* 220 K at ambient pressure. It corresponds to a transition from a low density liquid (LDL) to a high density liquid (HDL). In the case of bulk water it is assumed to imply a transition of the connective properties of the hydrogen-bonded network from an almost crystalline tetrahedral network (LDL) to disordered interconnecting network with a high degree of interstitial sites (HDL) [55]. In the case of interfacial water this transition occurs with the creation of patches of HDL water in the preexisting LDL phase and is combined with a strong increase of mobile water molecules [56]. This has also been held by some authors as the explanation for the dynamical transition of hydrated proteins [57, 58]. Therefore, at *T =* 220 K the dynamical variable is trapped in local minima requiring some energy to escape the trap set by the ordered deuterium-bonded network (*ϵ*_1_/_2,220K_ ≈ 2), while at *T* = 310 K the potential drastically softens and the thermal energy is enough to overcome the barriers (*ϵ*_1_/_2,310K_ ≈ 1).

The former example shows how the environment of the protein modifies the local landscape seen by the averaged self dynamical variable assuming largely incoherent scattering. Furthermore, it can provide a comparison of the landscapes of individual and collective dynamical variables, if polarized neutrons are used. The successful separation of both coherent and incoherent contributions at *T =* 310K indicates that in our sample, self and collective variables see a similar energy landscape, as indicated by *ϵ*_1_/_2_ falling within error bars for both incoherent and coherent scattering. The median energy *ϵ*_1_/_2_ required to escape a local minimum is of the order of one, meaning that Δ*E* is of the order of *k*_*B*_*T* . This is due to water being the main scatterer of the hydrated protein powder in this energy and *q*-ranges. However, such an analysis could reveal distinct dynamics if performed at slower time ranges and smaller *q*-ranges, corresponding to long-range motions within the protein.

### C. The coherent contribution to elastic scattering

Elastic scattering on proteins has been discussed for decades, often in relation with the dynamical transition at approximately *T =* 220 K [53, 54], where the slope of the MSD changes drastically. However, the temperature dependence of coherent elastic scattering of proteins and its impact on the dynamical transition remains, to our knowledge, a largely unexplored area. It has been shown that a dry powder of a deuterated protein undergoes a dynamical transition [59], seemingly due to the activation of collective modes implying heavy atoms. Furthermore, a fully deuterated powder of C-phycocyanin was studied by Bellissent-Funel *et al*., for which they calculated the coherent elastic structure factor at different temperatures in dry and in D_2_O hydrated states, showing a non-Gaussian behaviour of the ECSF and a drop of the Debye-Waller factor around the dynamical transition [36]. Otherwise, no comparison has been made of coherent versus incoherent elastic scattering, their fits following equation 27 are compared for *T* = 295 K. and to what extent both contribute to the amplitude and temperature dependence of the position fluctuations in a hydrated protein powder. As described in Materials and Methods (section III B 2), elastic fixed window scans were performed with similar resolutions on IN5 and LET, where incoherent and coherent scattering have been separately measured on LET. MSDs are calculated for both instruments in the range *q* ∈ [0.5, 2.2] Å ^−1^, using the Gaussian approximation given in eq. 27.

On Figure 4 a), where lines are a guide to the eyes, MSD_tot,IN5_ are calculated using IN5 non-polarised elastic data in the range *T* ∈ [14, 300] K (green triangular markers), following the usual assumption that the coherent component of elastic scattering is negligible. The ESF slightly differ from IN5 to LET: MSD_tot,LET_ are also calculated for comparison by summing spin flip and non-spin flip intensities on LET (orange circular markers). Therefore, 12 minutes scans are enough to estimate the MSDs on LET for *T >* 50K.

**FIG. 4.**
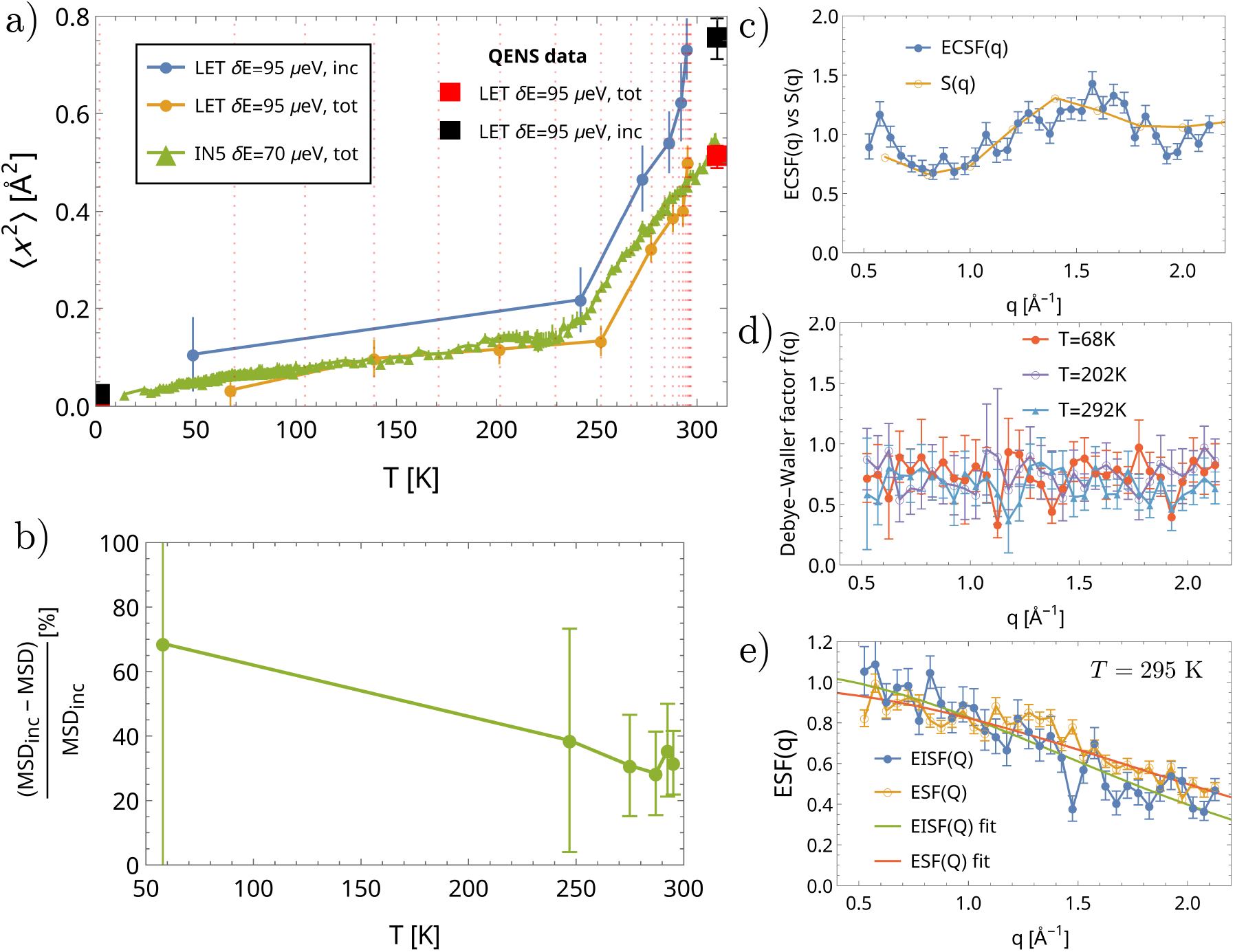
a) MSDs denoted ⟨*x*^2^⟩ are represented as a function of temperature *T*, for 14 K ≤ *T* ≤ 310 K. Lines are a guide for the reader. In green Δ markers, MSD_tot_ obtained with unpolarized neutrons on IN5 with energy resolution *δE* ≈ 77 *µ*eV (5 minutes elastic fixed window scans). In comparison, data obtained with polarized neutrons on LET with energy resolution *δE* ≈ 95 *µ*eV are represented with orange and blue ° markers, corresponding to total and incoherent scattering respectively (longer scan performed during sample heating, temperature sampling is shown by vertical red dotted lines). For both instruments, MSD_inc_ and MSD_tot_ are calculated using the *q* range [0.5, 2.2]Å^−1^ in the frame of the Gaussian approximation, Eq. 27. Elastic fixed window scans are corroborated by the model-dependent EISF (black □) and ESF (red □) calculated with LET data at *T =* 2 K and *T =* 300 K from Eq. 15. b) The relative error of incoherent with respect to total MSDs is displayed as a function of temperature *T* . c) The elastic coherent structure factor, blue curve, is compared to the static structure factor, orange curve. *S(q)* and ESCF(q) are found not to change significantly with temperature and therefore are averaged over all temperatures to improve the signal-to-noise ratio. Both are normalized to reach 1 for *q* → ∞, despite the reduced *q*-range. ; d) The Debye Waller factor defined in equation 29 is shown for three temperatures chosen in the range of study. e) EISF(*q*) and ESF(*q*) (filled and open markers respectively) and their fits following equation 27 are compared for T = 295 K.

However, the Gaussian approximation and more refined models for the calculation of MSDs [40, 41] require using EISF_m_ instead of ESF_m_, and is used under the assumption that collective dynamics are negligible. To the best of our knowledge, this has never been performed because of the limited access to polarized neutrons. In order to quantify the contamination of coherent elastic scattering in the estimation of MSDs, we calculated MSD_inc_ using the spin-flip intensity only (blue dots). Comparing MSD_tot_ and MSD_inc_, it appears that MSD_inc_ are significantly larger (≥ 30%) over the whole temperature range from *T =* 50K to *T =* 310 K, see Figure 4 b), especially above the dynamical transition. However, our temperature range does not permit to elaborate whether a shift of the dynamical transition occurs due to coherent scattering contamination.

To verify the precision of our findings, we also calculate MSD_inc_ and MSD_tot_ for *T =* 2K and *T =* 310K using the quasi-elastic scattering data integrated over the elastic peak. Scattering events were measured on longer times; therefore the signal to noise ratio is high compared to the fast elastic window scans. The MSD_inc_ are displayed in black squares, MSD_tot_ in red squares. It emphasizes the quality of results since the short time acquisitions align with long measurements: the discrepancy between MSD_inc_ and MSD_tot_ is indeed caused by coherent scattering contamination, which was already suggested to be non-negligible by Gabrys *et al*. [11].

To investigate why MSD_inc_ are systematically higher than MSD_tot_, we write down the coherent elastic intensity as the product of the static structure factor *S*(*q*) and the coherent Debye Waller factor *f* (*q*) following Bellissent-Funel *et al*. [36]:

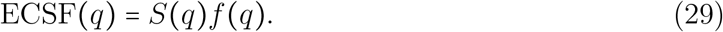

We evaluate *S(q)* for each temperature using the integrated coherent intensity on the full energy range (*E* ∈ [−1.5, 1.5] meV), in order to extract the Debye-Waller factor *f(q)*, see Figure 4 d). *f(q)* is almost constant over *q* for all temperatures, and therefore barely contributes to the monotonous decrease of ESF(*q*). Therefore, the structure of *S(q)* (commented further in the next section) is mostly responsible for the shape of ECSF(*q*). It is shown on Figure 4 c) which compares ECSF(*q*), averaged over temperature, to the static structure factor *S(q)*. We observe that, in agreement with the results of Bellissent-Funel et al. obtained with a hydrated deuterated C-phycocyanin [36], the structural information arises in the ECSF(*q*) around the main broad inter-atomic correlation peak *q =* 1.5 Å^−1^ which is typical for protein samples [60]. Since the ratio of coherent over total elastic scattering is found to barely depend on temperature and that neither *S(q)* or *f(q)* evolve with temperature, it explains that the error made on MSD_tot_ (Figure 4 b) is consistent over the whole temperature range. As a consequence, ESF(*q*) is contaminated mostly by the correlation peaks of ECSF(*q*) distorting the monotonous exponential decay of the incoherent component of elastic scattering. This explains that MSD_inc_ are systematically higher than MSD_tot_, as positive correlation peaks are present at larger *q*-ranges where the decay of EISF(*q*) is the most pronounced. Besides this structural modulation, coherent elastic scattering is almost a flat line that does not decay or increase much with *q*.

We foresee that this contamination might have more significant impact, for instance, in a parametric study comparing proteins with different amounts of hydration. In that case we expect *S(q)* to have different *q* and *T* dependencies due to the confined water structure and dynamics changing with the size of the hydration layer [60, 61]. Furthermore, the Debye-Waller factor could show more complex behaviour over *q* as observed in the case of Bellissent-Funel *et al*. in the C-phycocyanin protein [36], and contribute to the decay of the ESF. The comparison of different proteins will also suffer from S(q) depending on the secondary structure of the protein [60].

### D. Experimental and calculated *S*(*q*): a guide for polarized QENS experiments

Spectroscopic studies using polarized neutrons are very demanding: the flux is strongly reduced due to neutron absorption by the polarizer and analyser, and currently very few neutron spectrometers offer such a polarizing setup. Furthermore, proteins are incoherent scatterers and therefore require long time exposure to increase the signal to noise ratio of coherent scattering in the QENS region. As also discussed previously [21], the preparation of deuterated samples is tricky and requires precise monitoring of air composition during hydration.

For that reason, measuring the structure factor on polarized diffractometers prior to polarized QENS measurements would provide key information, such as the ratio of coherent scattering and its *q*-dependence, in order to further optimize the *q* and *ω* ranges required for QENS.

Figure 5 a) provides a set of measurements for *S(q)* performed on the deuterated GFP. Owing to the prevalence of elastic scattering in the hydrated protein, different instruments integrating from the elastic peak only (IN12 at ILL, −*E*_min_ = *E*_max_ = 0.5 meV) up to a large range of energies (D7 at ILL, *E*_min_ = −25 meV, *E*_max_ = 3.55 meV), and even at different temperatures (LET at ISIS), all provided the same *S*_coh_(*q)/S*_tot_ (*q)* ratio. This experimental result shows the reproducibility of the form factor over time. A first important observation is the presence of structure over the *q*-range used in the current study, which is the sign of non-negligible cross-correlation in coherent scattering. A second observation is the predominance of coherent scattering for *q* 05 Å^−1^., which is the range that usually goes with higher resolution QENS setups.

**FIG. 5.**
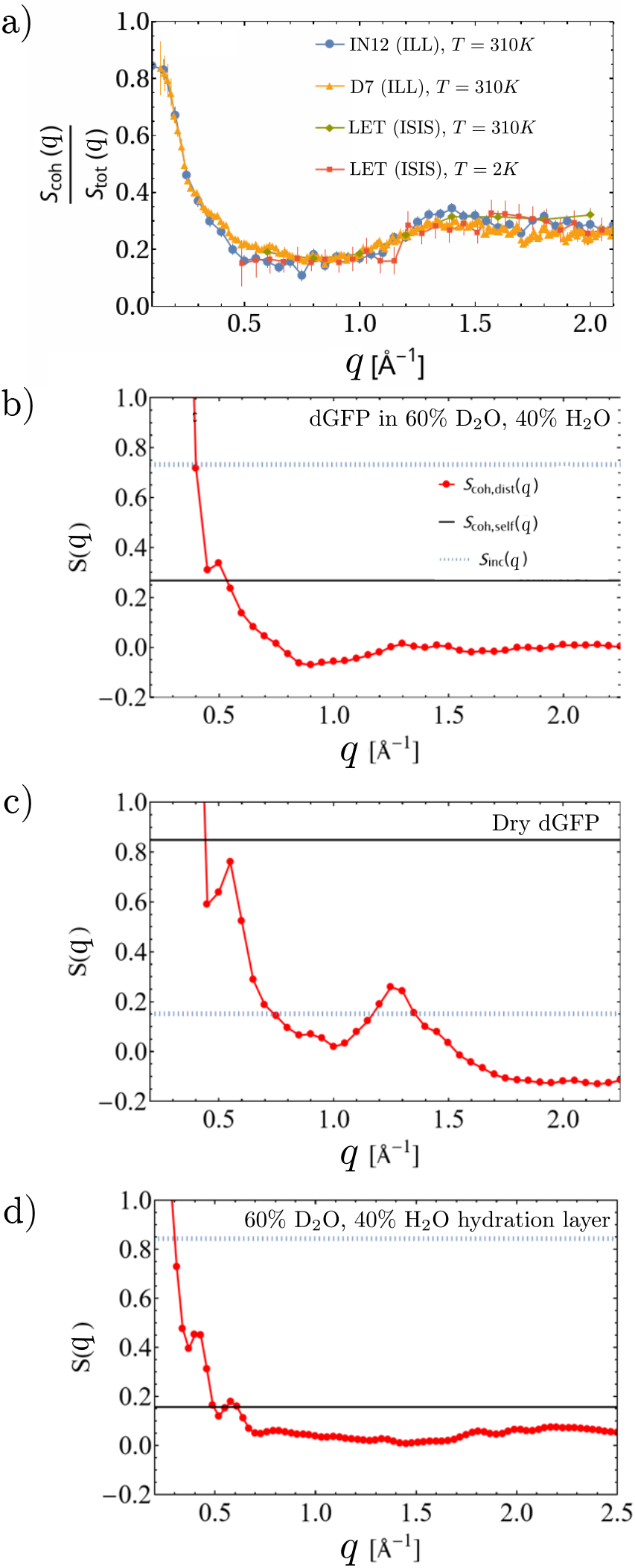
a) Experimental ratio of structure factors *S*_coh_(*q)/S*_tot_ (*q)* measured on D7 (diffractometer), IN12 (triple axis spectrometer) and LET (time of flight spectrometer), each showing different integration ranges. Further information is available in Ref. [21]. b-d) Calculations of the structure factor of a dGFP protein surrounded by a thin layer of 40% H_2_O and 60% D_2_O. b), c) and d) respectively correspond to (b) the hydrated dGFP, (c) the dry dGFP, (d) the hydration layer. *S(q)* and divided into its different components, see Eq. 31. The coherent distinct term *S*_coh, dist_ is depicted in red (structural term, *q* dependent), the coherent self term *S*_coh, self_ in black, and the incoherent self term *S*_inc, self_ in dotted blue lines.

To unravel which nuclei are responsible for the structural coherent scattering in the hydrated protein, exhaustive calculations of the structure factor of the GFP protein are performed. Assuming that the powder sample protein is isotropic, the static structure factor is given by,

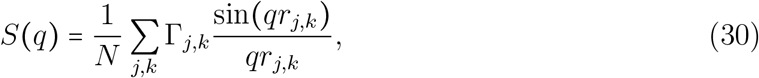

with *r*_*j,k*_ the scalar distance between nuclei *j* and *k*, and 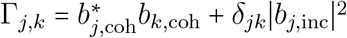. More details are available in Ref. [21]. It is important to keep in mind that coherent scattering is indeed the linear combination of a ‘distinct’ contribution, which arises from distinct nuclei, and a ‘self’ contribution which arises from correlations of the same nucleus and is therefore constant over *q* (since *r*_*j,j*_ *=* 0) as is the incoherent part. Both are weighted by the coherent scattering lengths. This is, then, formally expressed as,

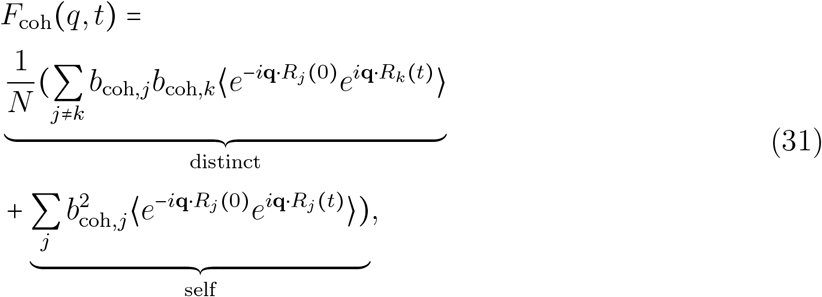

where *j* and *k* are distinct nuclei. Therefore, self-scattering in coherent QENS can be non-negligible, warranting a cautious use of the term ‘collective motions’ when referring to coherent inelastic scattering. The competition between the self and distinct terms is illustrated in the case of bulk D_2_O solvent in Ref. [62]. As a consequence, *S*_coh_(*q)* is also separated into self and distinct terms.

Calculations indeed provide a better understanding of the respective importance of self and distinct components in *S(q)* . First and foremost, calculations were performed for our specific sample constituted of the dry dGFP protein and its surrounding water (*h =* 0.4, 60% D_2_O and 40% H_2_O). In the dry protein, see Figure 5 c, the coherent self-component in black is dominating. In the water mixture only, see Figure 5 d, the lack of inter-nuclei correlations (red markers) and a low level of 20% coherent self-scattering (black line) makes it dominantly an incoherent scatterer. Therefore, the large mass ratio of water in the sample implies that eventually, the hydrated sample (Figure 5 b) is only ≈ 30% a coherent scatterer. Figure 6 in the appendix provides a detailed separation of *S*(*q*) into all atomic distinct and self terms for *q* > 05 Å^−1^. . It appears that the peak around *q* = 1.5Å^−1^ is mostly driven by C,O and N nuclei which compose the protein’s backbone. It is due to the *β*-barrel secondary structure that implies ordering of the carbon backbone. On the other end, correlations implying hydrogen isotopes are less structured and lower in intensity, as H and D nuclei are more randomly distributed in the side chains at *T =* 310 K.

**FIG. 6.**
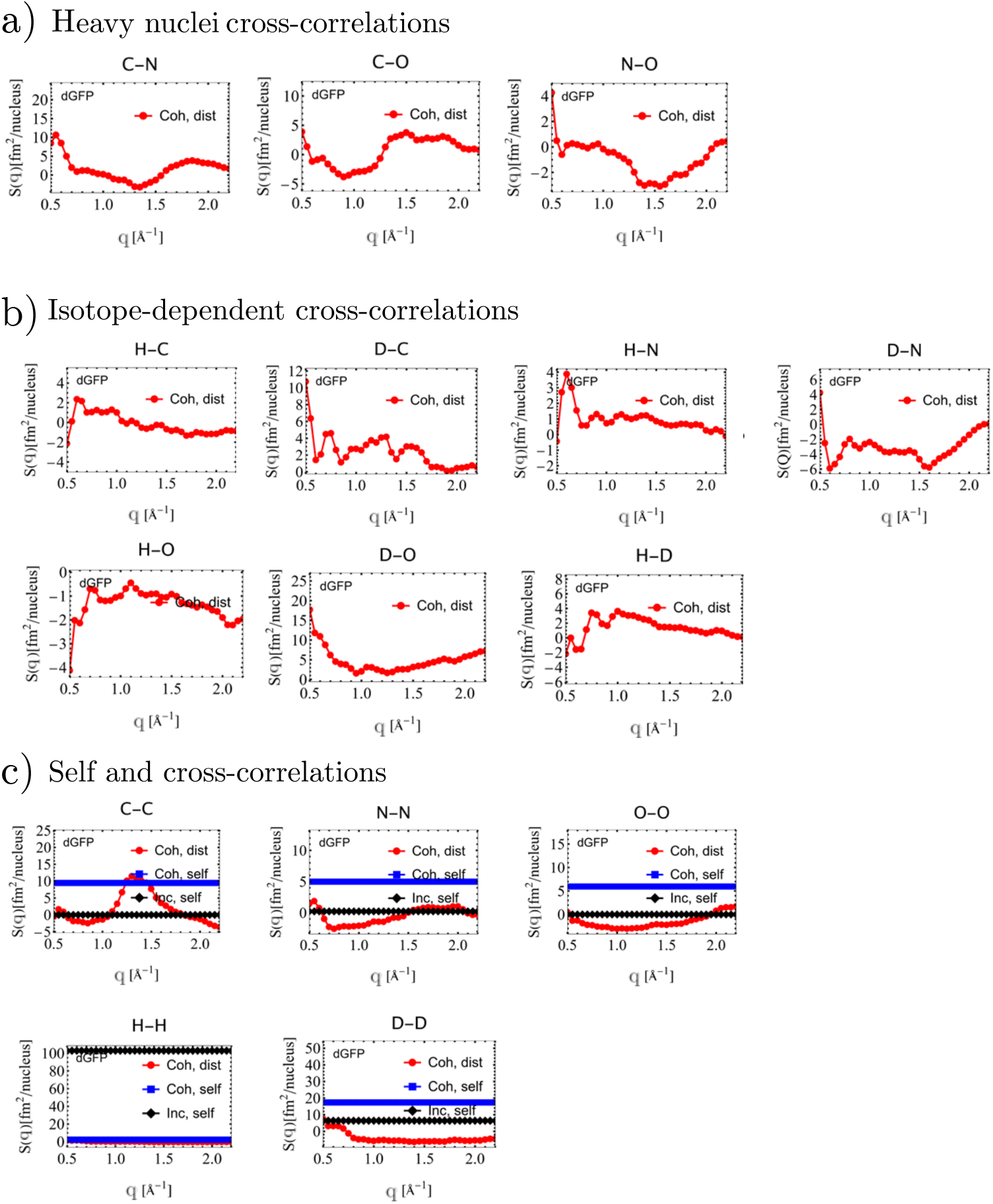
Incoherent self, coherent self, and distinct contributions to the structure factor *S(q)* of the dry GFP protein, calculated from wild-type deuterated GFP 6l26 PDB structure [75] assuming an isotropic sample and normalized per nucleus contribution. (a) Provides the distinct contributions from main heavy nuclei C,N and O, (b) provides the distinct contributions involving H and D nuclei, and (c) provides the distinct and self contributions from similar nuclei types.

Although the *q*-range presently studied favours the contrast of the backbone, the time-window restricted by the energy range available on LET emphasizes the very fast dynamics of water with respect to slow backbone dynamics. In the analysis of the QENS data, see Figure 2, it accounts for the structure visible in the ESF around *q =* 1.3 Å^−1^ (immobile protein backbone) counterbalanced by fast dynamics of water (*τ* parameter).

To extend this analysis further, a complete description of the distinct (namely, purely collective) contributions to *S*_coh_(*q)* for a fully deuterated protein in 100% D_2_O is provided on Figure 7 in the appendix. It is the ideal case study to extract collective dynamics of hydrated protein powders, even with an unpolarized beam of neutrons [63, 64]. Figure 7 a) displays all distinct components constituting *S(q)* for a GFP protein hydrated with *h =* 0.4 g D_2_O/g protein. It appears that D-O correlations arising from D_2_O dominate the structural intensity for *q* < 1 Å^−1^. Above that range, correlations stem from both the protein’s backbone and deuterated water. It predicts that yielding collective information from the protein only, in the present *q*-range, is non trivial due to the presence of water.

**FIG. 7.**
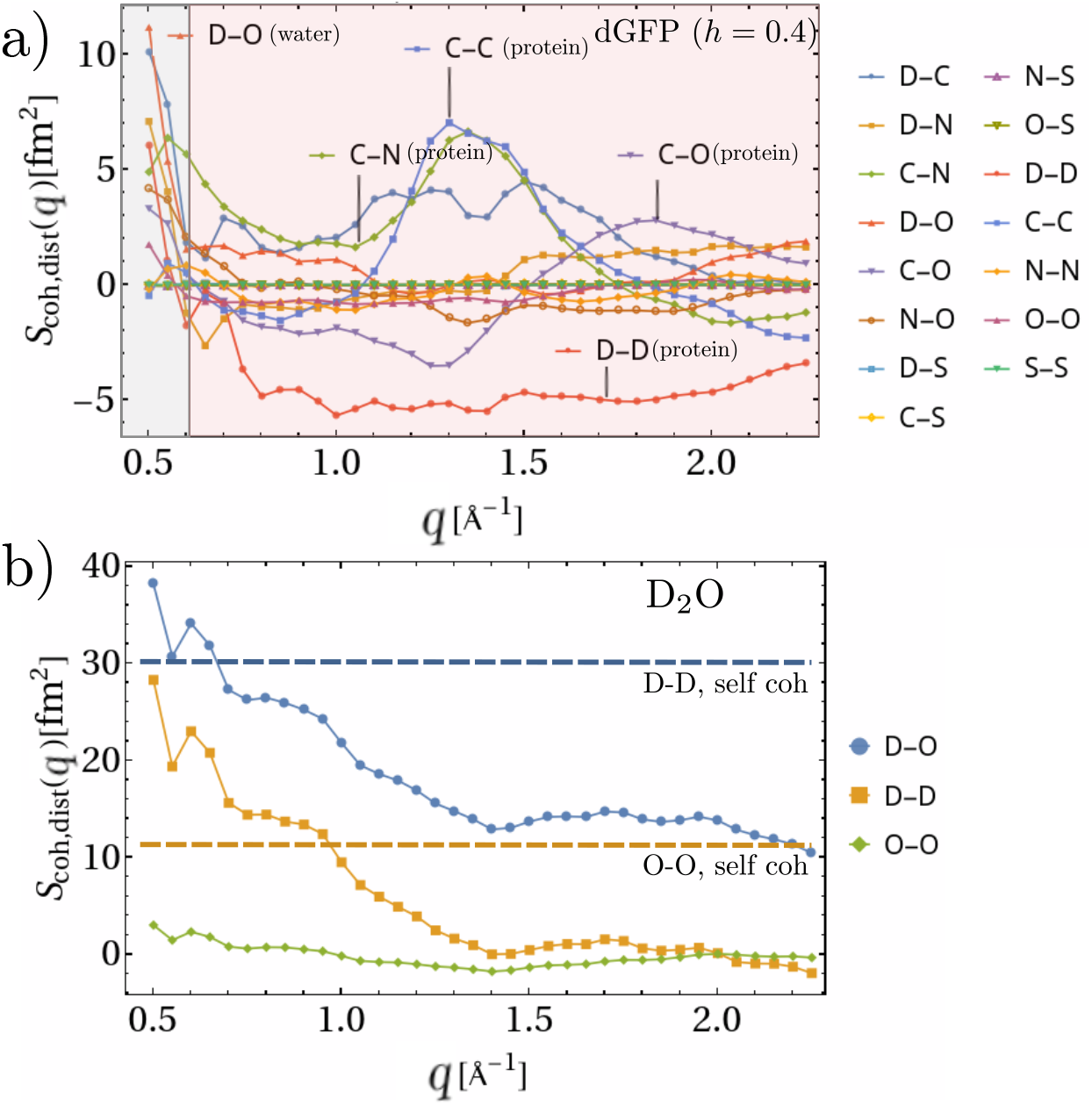
Calculated distinct coherent contributions of all pairs of nuclei (joined markers) for (a) a dGFP protein in *h =* 0.4 D_2_O and (b) the D_2_O hydration water only. in b), self coherent contributions are depicted using dashed lines.

A highlight on D_2_O hydration layer is provided in Figure 7 b. Self-scattering represents more than 70% of coherent scattering for *q* > 0. 5 Å^−1^. It is indeed of upmost interest to compare with the study of streptavidin protein in solution by Sarter et al [4], which shows that water overtakes the dynamics for *q* > 1.15 Å^−1^. Our work suggests that coherent self-scattering of water might be responsible for most of the contamination, rather than collective dynamics of water.

Another comparative study is that of Gaspar et al [60]. They proved experimentally that both hydration and bulk water strongly affect *S*(*q*). Hydrating or solubilizing the protein have opposed effects on *S*(*q*) for *q* < 0. 5Å^−1^ . While hydration increases the coherent contribution, increasing bulk D_2_O concentration in solubilized proteins lowers the coherent contribution. Indeed, the *q*-dependence of *S(q)* for a dry globular protein is expected to rely on secondary structure merely through changes of *C ™ x* peak positions, and does not depend much on isotope exchange. However, water is paramount in both tuning the intensity of coherent scattering and its *q*-dependent structure. Since lim_*q→*0_ *S(q) = ρk*_*B*_*Tχ*_*T*_ with *χ*_*T*_ the isothermal compressibility, it is indeed expected that a protein powder has higher *S(q)* for *q <* 0.5 Å^−1^ with respect to a protein solution.

A clear output from both experiments and calculations is the importance of isotope exchange and water content to tune the intensity of coherent scattering. Furthermore, self-scattering (either incoherent or coherent) is dominant in the case of hydrated proteins.

Therefore, it renders difficult the study of ‘collective dynamics’ in proteins without proper evaluation of relative coherent to incoherent intensities, and the knowledge of self and distinct components within coherent scattering. Even once calculations or diffraction performed, it is not even certain that *S(q)* reflects the effective relative intensity of nuclei in the range of energy under study as shown in our case, and cannot replace molecular dynamics simulations or normal mode analysis simulations [64]. Therefore, estimating coherent fast dynamics in proteins is currently a challenge. Using neutron spin echo with lower *q* and *ω* ranges could also bridge the gap to make use of the important coherent scattering in the SANS region.

## V. CONCLUSION AND DISCUSSION

We have studied elastic and quasi-elastic neutron scattering in a deuterated protein powder with two extremely similar instruments, respectively IN5 and LET, belonging to the same category of time-of-flight neutron spectrometers. However, only LET is equipped with a complete set-up to perform uniaxial polarization analysis, enabling unambiguous separation of incoherent and coherent scattering. We show that the use of a minimalistic model accounting for fractional Brownian dynamics enables comparison of self and collective dynamics in the context of complex samples showing self-similar dynamics over large time-ranges. It also facilitates comparison between different instruments. The model therefore provides a framework to study coherent scattering in complex biological systems with a large range of diffusion timescales, which so far required purely deuterated samples and strong assumptions on the relative importance of coherent cross-sections. Using a deuterated GFP in mixed solvent due to quasi-instantaneous H → D exchange in the hydration layer (more information in Ref. [21]), we provide comparison of scattering functions and parameters of IN5 and LET. It is followed by comparison of dynamical parameters obtained for incoherent and coherent scattering and their interpretation in terms of energy landscape, where we capture a significant change around the dynamical transition corresponding to hydration water. Then, we show that the ECSF closely follows the structural modulations of the static structure factor *S(q)* . It implies that the exponential decay of EISF as a function of *q* is distorted by the presence of correlation peaks, especially at *q* ≈ 1.5Å^−1^. As a consequence, MSDs obtained with the ESF using the fixed window scan method with unpolarized neutrons are significantly under-estimated (≈ 30% in our study) over the whole temperature range from *T =* 2 *K* to *T =* 300 *K*. This could have substantial impact on the comparison across protein samples, especially when different hydration rates are used, since it largely impacts the shape of the structure factor in the *q*-ranges studied in the elastic fixed window scan method. Finally, biological samples should always be in hydrated state. As reported earlier in Refs. [4, 21], water coherent scattering plays an important role in the total scattering of the hydrated or solubilized protein. Hence, we propose that diffraction studies performed along with structure factor calculations on the *q*-range of study should help decipher all contributions from the protein and its hydration layer, and therefore provide a good insight into the relative importance of the nuclei before a QENS experiment. It should also provide an estimation of self and distinct components of the scattering function, informing whether coherent scattering indeed provides insight into collective dynamics, or whether it contains significant self-scattering contributions. In that regard, we expect that both the heavy protein backbone and the close protein hydration layer contain non-negligible self-scattering in their coherent scattering term, questioning the use of coherent scattering for collective dynamics studies with the current accessible techniques and *q*-ranges. However, *S(q)* might not be representative of QENS self and distinct components if the sample is highly dynamic in the considered energy range, or if different nuclei follow different dynamics as it is the case for our GFP sample.

Hence, using polarized neutrons for complex non-magnetic samples merits investigation. Recently, there has been an increased interest for experimental investigation of collective dynamic scattering in proteins [4, 21], ionic liquids [65, 66], solvents [67–70], polyelectrolytes [71] or even small drugs [72]. From these recent results emerge the strong potential of polarized neutrons correlated to molecular dynamics as a tool to understand better the collective dynamics of solvents and small molecules in solution. For instance, Arbe et al. [67] used the comparison of incoherent and coherent scattering to explain the universal presence of a structural mode in self-dynamics of fluids showing different types of interactions, and Morbidini et al. [72] explored the mesoscopic origins behind the non-linear phase diagrams of binary mixtures. However, the size and amourphous character of proteins imply that using polarized neutron scattering to study collective dynamics will require additional information from molecular dynamics simulations, and will not simply rely on model fitting. Concerning the use of polarized neutrons to estimate the impact of coherent scattering in the total spectrum, it will require more systematic studies and varying protein structures in order to assess the necessity of using polarized neutrons in conventional QENS studies of hydrogen-rich proteins. However, our work points out an important scattering of the hydration layer, which will significantly depend on the hydration level and on the *q*-range under study, suggesting that drawing general conclusions will not be possible. Still, we observe that the hydration layer of a powder protein does not affect dynamical parameters enough to justify the use of polarized neutrons for coherent contamination issues. In particular, the precision on parameters gained by using polarized neutron scattering on our protein sample where incoherent scattering dominates (80% of total scattering, see Figure 5) does not compensate significantly the inter-instrumental error when measuring the same sample on two similar instruments, as can be seen by comparing blue, gray and black markers on Figure 2. It means that performing experiments on hydrogen-rich samples with high incoherent cross-section on polarized instruments instead of conventional non-polarized instruments might not be relevant on all *q* scales. Nevertheless, Sarter et al [4] suggest otherwise for hydrogen-rich protein solutions, where they draw a clear *q*-range above about *q =* 1 Å^−1^ where coherent scattering is supposed to become more and more predominant. This q-cut-off value is, however, resolution dependent, and results from Gaspar *et al*. [60] also suggest that this threshold will be D_2_O-concentration dependent. In the case of a protein powder, our calculations for *S(q)* (Figure 5) and QENS measurements suggest that large *q* ranges will see dominance of water due to its fast dynamics at large *q*-ranges, and probably also at low *q*-ranges due to the increasing *S*_coh_(*q*)/*S*(*q*) ratio.

Indeed, different factors justify the use of polarized neutrons only when the study requires it. To begin with, the counting-rate of neutrons is reduced by a factor of about 10. This is due to 60% of flux lost due to the polarization process, then multiplied by the loss of neutrons due to the spin-analyser transmission. It makes extensive studies difficult to perform, such as monitoring different environmental conditions for the samples. Then, polarized QENS is currently accessible up to the pico-second scale: although a 10*µ*eV resolution is achievable on LET, its low flux requires long measurements in order to have a sufficient coherent signal to noise ratio in the QENS region. Finally, not all instruments are adapted for the installation of a polarization set-up. For the time being, the possibility to use neutron polarization analysis for high-resolution QENS is scarce, although more instruments are being deployed worldwide [73, 74].

We have shown, however, that polarization analysis proves to be a necessary tool to investigate unexplored topics of protein dynamics such as coherent elastic scattering, or the comparison of individual and collective self-dynamics at short times inside a protein, or the impact of secondary structure on collective motions inside the protein. In this perspective, we offer the following guidance: it is important to study proteins with variable secondary structures or functions to investigate the effect of long range inter-domain motions, the impact of the native structure, the presence of disordered regions, etc. The quality of the deuteration process appears to be of paramount importance [21] and moreover, the development of theoretical models to account for local collective motions is pivotal in order to make sense of polarized neutron scattering experiments. In order to investigate lower *q*-ranges corresponding to more cooperative scales, combining these studies with the spin-echo technique using deuterated samples could prove to be effective.

## AUTHOR CONTRIBUTIONS

Conceptualization, J.P.; methodology, A.N.,J.P.; software, A.N.; validation, A.N., J.O., R.S.; investigation, A.N., J.P., J.O., R.S.; resources, J.P., J.O., R.S.; data curation, A.N., J.O., R.S.; writing—original draft preparation, A.N.; writing—review and editing, J.P, J.O, R.S; visualization, A.N.; supervision, J.P.; project administration, J.P.; funding acquisition, J.P. All authors have read and agreed to the published version of the manuscript.

## DATA AVAILABILITY

Instrumental data are openly available under the following DOIs: 10.5286/ISIS.E.RB2220225 (LET), 10.5291/ILL-DATA.DIR-173 (ILL). Further data or software are available upon reasonable request.

## ACKNOWLEDGMENTS

The authors thank G. Kneller for continuous discussions and help with this topic. This research was funded by Mission pour les Initiatives Transverses et Interdisciplinaires du CNRS, BioQuant, attributed to J. Peters. We acknowledge as well the PhD grant attributed to A. Nidriche, delivered by Ministère de l’éducation nationale. We thank Institut Laue-Langevin, Grenoble, France, for beamtime allowance on IN5 instruments (10.5291/ILL-DATA.DIR-173), as well as ISIS neutron facility for beam time allowance on LET (10.5286/ISIS.E.RB2220225-1).

**Appendix: Extensive description of cross-correlations of the scattering function of a dry protein, and a protein hydrated in D**_2_**O (***h* = 0.4).

